# OKseqHMM: a genome-wide replication fork directionality analysis toolkit

**DOI:** 10.1101/2022.01.12.476022

**Authors:** Yaqun Liu, Xia Wu, Yves D’aubenton-Carafa, Claude Thermes, Chun-Long Chen

## Abstract

**Motivation:** During each cell division, tens of thousands of DNA replication origins are coordinately activated to ensure the complete duplication of the entire human genome. However, the progression of replication forks can be challenged by numerous factors. One such factor is transcription-replication conflicts (TRC), which can either be co-directional or head-on with the latter being revealed as more dangerous for genome integrity.

**Results:** In order to study the direction of replication fork movement and TRC, we developed a bioinformatics tool, called OKseqHMM, to directly measure the genome-wide replication fork directionality (RFD) as well as replication initiation and termination from data obtained by Okazaki fragment sequencing (OK-Seq) and related techniques.

**Availability and Implementation:** We have gathered and analyzed OK-seq data from a large number of organisms including yeast, mouse and human, to generate high-quality RFD profiles and determine initiation zones and termination zones by using Hidden Markov Model (HMM) algorithm (all tools and data are available at https://github.com/CL-CHEN-Lab/OK-Seq). In addition, we have extended our analysis to data obtained by related techniques, such as eSPAN and TrAEL-seq, which also contain RFD information. Our works, therefore, provide an important tool and resource for the community to further study TRC and genome instability, in a wide range of cell line models and growth conditions, which is of prime importance for human health.

**Contact:** Chun-Long Chen (Institut Curie)

**Category:** **Genome analysis**

## 1. Introduction

The faithful transmission of genetic information to daughter cells is crucial for maintaining genome stability. In humans, at each cell division, tens of thousands of replication origins need to be coordinately activated to ensure the complete duplication of >6 billion base pairs (bp) of the human genome. However, the DNA replication program is routinely exposed to endogenous and exogenous stresses, which play an important role in many human diseases. In particular, replication stress-induced genome alterations can represent an important early cause of cancer (Gnan *et al.*, 2020).

The progression of replication forks can be challenged by numerous factors. One such factor is transcription-replication conflicts (TRC) since the replication and transcription machineries share the same DNA template. TRC can either be co-directional (CD) or head-on (HO), and the latter has been revealed as more detrimental for genome integrity (Hamperl *et al.*, 2017). Previous bioinformatics analyses have revealed that, in large numbers of genomes from bacteria (Merrikh *et al.*, 2017) to human (Chen *et al.*, 2011; Huvet *et al.*, 2007), most genes are co-directionally oriented with replication forks to avoid the more deleterious HO TRC. Recently, a new method to directly measure the genome-wide replication fork directionality (RFD) along the human genome by sequencing of Okazaki fragments (OK-Seq) (Petryk *et al.*, 2016), which are present only on the lagging replicating strand, allows quantitatively analyzing and accurately detecting replication initiation and termination. The analysis of OK-seq data of human cells has also demonstrated a significant co-direction of replication fork progression with gene transcription within active genes (Petryk *et al.*, 2016).

More and more techniques are now being developed, for instance, Pu-seq (Daigaku *et al.*, 2015), eSPAN (Li *et al.*, 2020), SCAR-seq (Petryk *et al.*, 2018), GLOE-seq (Sriramachandran *et al.*, 2020) and TrAEL-seq (Kara *et al.*, 2021), which also provide genome-wide RFD information. Moreover, in recent years, strong evidence shows that replication- and transcription-related mutational asymmetries are widespread across cancer development (Haradhvala *et al.*, 2016). Especially APOBEC-associated mutations (also called APOBEC mutation signatures) in humans are represented in up to 15% of all sequenced tumors and contribute to 50% of all mutations in many tumors. APOBEC-associated mutations preferentially occur on the lagging-strand template during DNA replication, and are also highly associated with mismatch repair and transcription-coupled damage repair in cancer (Cortez *et al.*, 2019; Hoopes *et al.*, 2016; Shi *et al.*, 2019; M. J. Shi *et al.*, 2020; Mas-Ponte and Supek, 2020). Furthermore, N6-methyladenosine (m6A) modifications have been considered as one of the most prevalent internal modifications in mammalian mRNAs and the abnormal m6A modification caused by m6A modulators, e.g., methyltransferase-like 3 (METTL3), is a common feature of various tumors (Y. Shi *et al.*, 2020; Wang *et al.*, 2020; Huang *et al.*, 2020). Evidence has shown that METTL3 and m6A could promote homologous recombination-mediated repair of double-strand breaks (DSBs) by modulating DNA-RNA Hybrid (R-loops) accumulation (Zhang *et al.*, 2020). Importantly, R-loops have been recently shown, by others and us, to be frequently accumulated at transcription termination sites of actively transcribed genes displaying high HO TRCs (Promonet *et al.*, 2020; Liu *et al.*, 2021). Therefore, systematically unveiling the genome-wide DNA replication panorama is essential for human health.

Despite its importance, to date, there is no published available tool to analyze RFD data and to determine the replication initiation and termination positions genome-wide, although several methods have been previously described for OK-seq data analysis, such as using the Hidden Markov Model (HMM) to analyze human OK-seq data (Petryk *et al.*, 2016) or the origin efficiency metric (OEM) to analyze yeast OK-seq data (McGuffee *et al.*, 2013). It is, therefore, important to have a uniform framework of OK-seq data (and related data) analyses. Here, we developed a bioinformatics toolkit, called OKseqHMM, to directly obtain the high-resolution RFD profile genome-wide. Besides the fork direction, the toolkit also deciphers the information of replication initiation/termination zones using an algorithm based on HMM, calculates the OEM to visualize the transition of RFD profile at multiple scales, and finally generates the average metagene profiles and heatmaps to provide RFD/OEM distributions along the regions of interest (Fig. 1). We have gathered a large number of published available OK-seq data (13 in total) from *S. cerevisiae*, mouse and human cells, and successfully obtained the high-resolutive (~1 kb for mouse and human cells and ~50 bp for yeast) RFD profiles and the accurate calling of corresponding replication initiation and termination zones genome-wide.

**Figure 1.**
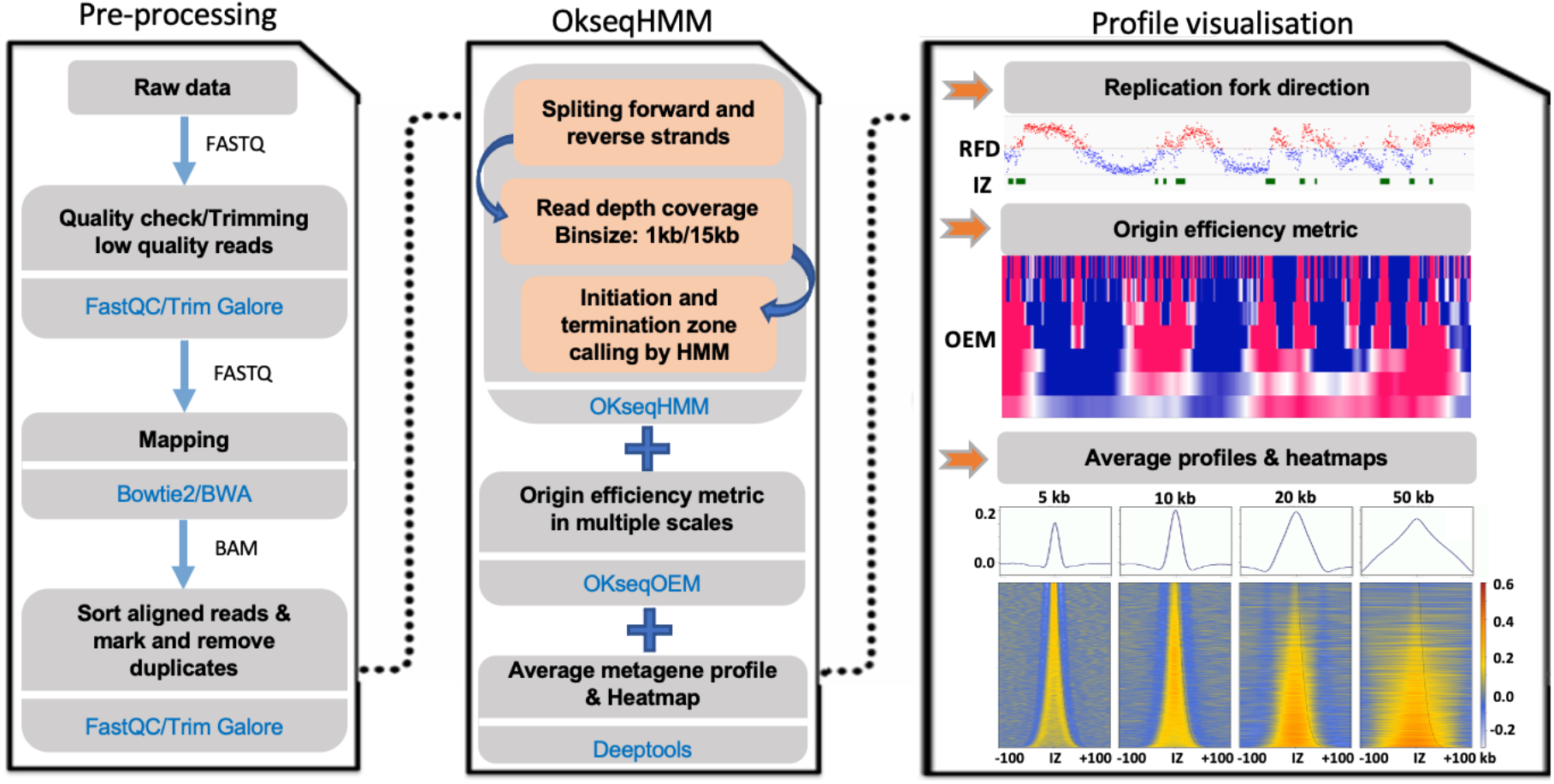
Schematic presentation of data analysis pipeline of OKseqHMM toolkit. The raw sequencing data can be pre-processed into aligned files by corresponding bioinformatics tools indicated in blue (left panel). The middle panel shows the major functions of the OKseqHMM toolkit. The first function of OKseqHMM checks the input aligned bam files to determine whether they are single- or paired-end sequencing data, then automatically splits the reads into Watson and Crick strands and calculates the replication fork directionality (RFD). By default, the calculation is performed within 1 kb adjacent windows (recommended for human cells) and then smoothed into 15 kb sliding windows with 1 kb step. These parameters can be easily adjusted based on the nature of the data. Different replication features, i.e., initiation zones (IZ), two intermediate states and termination zones, are predicted based on an HMM algorithm (See Implementation for detail). The second function (OKseqOEM) uses the reads on Watson and Crick strands to generate origin efficiency metrics (OEMs) at multiple scales to visualize the RFD transition. And the last function allows users to generate an average metagene profile and heatmap to analyze distributions of RFD and OEM around the genes/regions of interest. Results can be visualized in the genomic visualization browsers (such as IGV) as shown in the right panel.

## 2. Implementation and availability

OKseqHMM toolkit is an R package for profiling OK-Seq data to study the genome-wide replication program. This R package contains multi-functions and is served for analyzing OK-Seq data from the original mapping bam file(s) to count matrices, RFD calculation, initiation/termination zone calling and average metagene profiles/heatmaps. The package is available at https://github.com/CL-CHEN-Lab/OK-Seq.

### 2.1 Function OKseqHMM

This function transforms OK-seq data into RFD profiles for a primary visualization (e.g., with the genomic visualization browsers, such as IGV), then, it can accurately identify replication initiation zones (IZs, upward transitions on RFD profile), termination zones (TZs, downward transitions on RFD profile) and also the intermediate states (flat RFD profile) along the genome by using the HMM.

For each window, RFD was computed as follows:

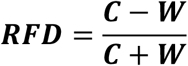

where ***C*** and ***W*** correspond to the number of reads mapped on the Crick and Watson strands, which reveal, respectively, the proportions of rightward- and leftward- moving forks within each window (e.g., 1 kb window was used for OK-seq data of human cells). Since the total amount of replication on both strands should be constant across the genome, we normalized the difference between the two strands by the total read count to account for the variations in read-depth due to copy number, sequence bias and so on. RFD ranges from −1 (100% leftward-moving forks) to +1 (100% rightward-moving forks), and 0 means equal proportions of leftward- and rightward-moving forks. Data obtained from biological replicates produced RFD profiles that strongly correlated to each other, for HeLa cells, Pearson R=0.92, p<10^−15^ (t-test) and for GM06990 cells, R=0.93, p<10^−15^. Similar correlations were observed between RFD profiles with EdU or EdC labeling (Petryk *et al.*, 2016).

As in (Petryk *et al.*, 2016), a four-state HMM was used in OKseqHMM to detect within the RFD profiles the ascending (AS), descending (DS) and flat (FS) segments representing regions of predominant initiation (’Up’ state), predominant termination (‘Down’ state) and constant RFD (’Flat1’ and ‘Flat2’ states) (Fig. 2A). In the HMM segmentation process, the RFD values were computed within 15 kb (for human OK-seq data) sliding windows (by default, stepped by 1 kb across the autosomes). The HMM used the Δ*RFD* values between adjacent windows, in which 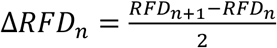 for the window *n*. By default, windows with <30 reads on both strands were masked. The Δ*RFD* values (also between −1 and 1) were divided into five quantiles and the HMM package of R (http://www.r-project.org/) was used to perform the HMM prediction with probabilities of transition and emission, which are manually defined by the training dataset (Fig. 2B). The same transition and emission probabilities used in our previous study (Petryk *et al.*, 2016) were set as default values and used in all OK-seq data analyses in the current study. The choice of a 15 kb sliding window is based on a compromise between spatial resolution and reproducibility of AS detection among biological replicates. Finally, the efficiency of the detected AS was estimated as:

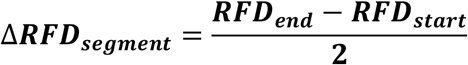

where ***RFD*_*start*_** and ***RFD*_*end*_** correspond, respectively, to the RFD values computed in 5 kb windows around the left and right extremities of each segment.

**Figure 2.**
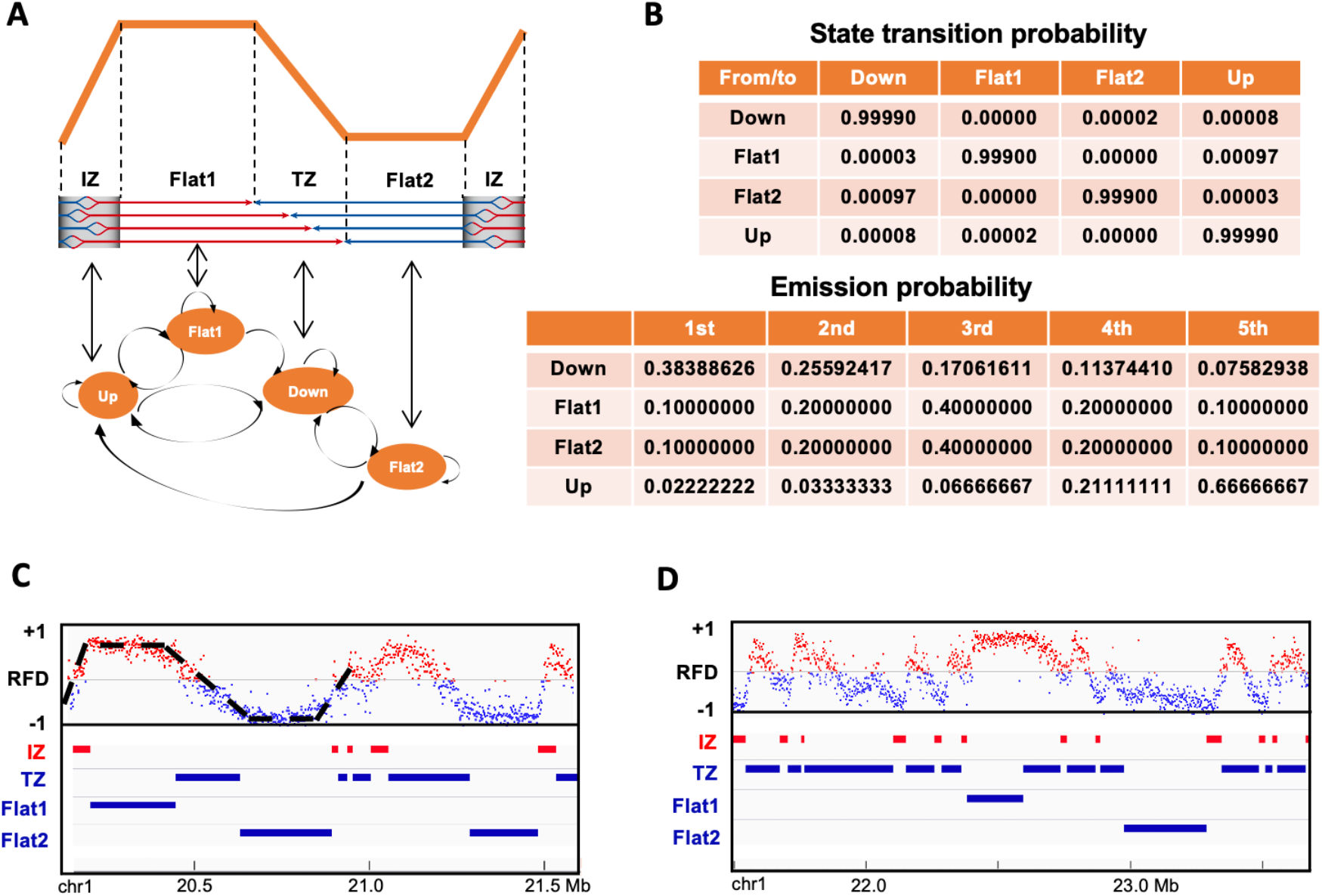
Schematic presentation of HMM algorithm for initiation and termination zone detection. **(A)** A 4-state HMM model used in the segmentation process: Up, regions of predominant initiation (IZ); Down, regions of predominant termination (TZ); Flat1 and Flat2, two intermediate transition states. **(B)** Transition and emission probabilities. **(C)** and **(D)** Examples of RFD profile of HeLa cells from chromosome 1 with the corresponding IZs, TZs and 2 Flat states identified by OKseqHMM. Each point on the RFD profile gives the RFD value calculated within each 1 kb adjacent window, and the windows with positive and negative RFD values are shown in red and blue, respectively.

### 2.2 Function OKseqOEM

For a further investigation of origin efficiency (i.e., Δ*RFD*), we provide here a second function to visualize it directly at multiple scales.

As defined in the previous publication for yeast OK-seq data analysis ^23^, the density of Okazaki fragments on the Watson and Crick strands are compared within 4 fixed-size sliding bins, which are strand-specific 10 kb quadrant values to calculate an Origin Efficiency Metric (OEM), computed as 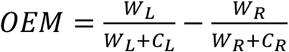 (*W*_*L*_ and *W*_*R*_ measure, respectively, the read density in the left and right quadrants on the Watson strand, while *C*_*L*_ and *C*_*R*_ refer to the density on the Crick strand), ranging from −1 to 1 for each base in the genome. Maximal values in the OEM scores represent replication origins, while the minimal ones are considered as regions of replication termination. In addition, the different amplitudes of positive OEMs (from 0 to 1) are referred to as origin-firing efficiency; and the degree of termination at each position can be measured from 0 to −1.

Here, we further extend OEM calculation within a fixed window size into multiple-scales to better fit OK-seq data analysis of other organisms, such as human cells.

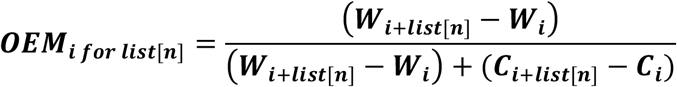

Where ***list[n]*** can be defined by users as a list of windows (e.g. [1,10, 20, 50, 100]), ***i*** is from 1 to the total length of the data – *list[n]*. ***C*** and ***W*** correspond to the number of reads mapped on, respectively, the Crick and Watson strands within corresponding windows.

Using the two bam files of reads within, respectively, Watson and Crick strands generated by the previous OKseqHMM function and the annotation coordinates, the function OKseqOEM can automatically calculate the OEM profiles at a series of defined scales (e.g., from 1 kb to 1 Mb for human cells), which allows us to directly visualize the transition states of replication and also to validate the IZs identified by OKseqHMM then to double-check the size and boundary of IZs.

### 2.3 Average metagene profile/heatmap

To analyze RFD distributions around and/or among the genomic regions of interest, such as the identified IZs or the annotated genes, we developed an additional module for the metadata analysis. With the gene coordinates (or IZs) together with the RFD and/or OEM big wiggle files generated from OKseqHMM and/or OKseqOEM functions, we can easily obtain the corresponding profiles/heatmaps by using the computeMatrix and plotProfile/plotHeatmap functions of deepTools (https://deeptools.readthedocs.io/en/develop/index.html) (Ramírez,*et al.* 2016) via defining the genomic distances of interest for the upstream and downstream borders.

## 3 Performance

### 3.1 Genome-wide replication fork directionality and origin detection in yeast

To evaluate the performance, we first applied our tool to the available yeast OK-seq data (Hennion *et al.*, 2020). OKseqHMM was successfully applied to the yeast OK-seq data to generate the RFD profile at a fine resolution (50 bp), the OEM profiles at different scales (from 50 bp to 25 kb) and a precise IZ/Origin calling (Fig. 3A). About 350 IZs were identified by OKseqHMM, which range from 0.5 kb to 5.5 kb with an average length of 1.5 kb (Fig. 3B, Table 1). To check the accuracy of IZ calling results, we compared OK-seq IZs with the known yeast origins, i.e. autonomously replicating sequence (ARS) from OriDB 2.1.0 (Siow *et al.*, 2012), and up to 80% of our detected IZs were found at ≤ 1 kb distance from a known ARS (Fig. 3C and D).

**Table 1.** All OK-seq data analyzed by OKseqHMM.

| Name | Cell type | Replicate | Initiation zones |  | Termination zones |  | Accession number<br>(Reference) |
| --- | --- | --- | --- | --- | --- | --- | --- |
| | | | Number | Size (kb)<br>Mean ± SD | Number | Size (kb)<br>Mean ± SD | |
| BL79 | Burkitt's lymphoma | | 7798 | 29±18 | 7791 | 211±244 | ENA: PRJEB25180<br>(Wu *et al.*, 2018) |
| GM06990 | Lymphoblastoid cells | 2* | 5684 | 33±19 | 5715 | 182±166 | SRA: SRP065949<br>(Petryk *et al.*, 2016) |
| HeLa MRL2 | Epithelial cell of adenocarcinoma | 2* | 9836 | 31±18 | 9441 | 141±144 | SRA: SRP065949<br>(Petryk *et al.*, 2016) |
| HeLa S3 | Epithelial cell of adenocarcinoma | | 9089 | 32±19 | 9084 | 223±245 | (Current study) |
| IARC385 | B lymphocytes from Burkitt's lymphoma | | 4465 | 36±19 | 4455 | 125±164 | ENA: PRJEB25180<br>(Wu *et al.*, 2018) |
| IB118 | Leiomyosarcoma | | 3645 | 26±16 | 3640 | 428±440 | ENA: PRJEB25180<br>(Wu *et al.*, 2018) |
| IMR90 | Fibroblast | | 12482 | 26±17 | 12468 | 151±147 | ENA: PRJEB25180<br>(Wu *et al.*, 2018) |
| K562 | Late-stage chronic myeloid leukemia | | 6982 | 28±15 | 6967 | 136±158 | ENA: PRJEB25180<br>(Wu *et al.*, 2018) |
| mESC E14 | Mouse embryonic stem cells | | 3370 | 27±14 | 3347 | 483±554 | GEO: GSE142996<br>(Li *et al.*, 2020) |
| Raji | Burkitt's lymphoma | | 8096 | 29±16 | 8080 | 143±135 | ENA: PRJEB25180<br>(Wu *et al.*, 2018) |
| TF1 | BCR-ABL negative cell line from erythroblast | | 8377 | 27±17 | 8371 | 196±193 | ENA: PRJEB25180<br>(Wu *et al.*, 2018) |
| TLSE19 | Leiomyosarcoma | | 10500 | 27±17 | 10492 | 146±144 | ENA: PRJEB25180<br>(Wu *et al.*, 2018) |
| Yeast | *S. cerevisiae* | 2* | 348 | 1.5±0.7 | 787 | 14±13 | ENA: PRJEB36782<br>(Hennion *et al.*, 2020) |

\* If data of biological replicates are available, the profiles obtained with the combined data are used in the figures, and only the segments (i.e. IZs and TZs) reproducibly identified in both biological replicates were retained.

**Figure 3.**
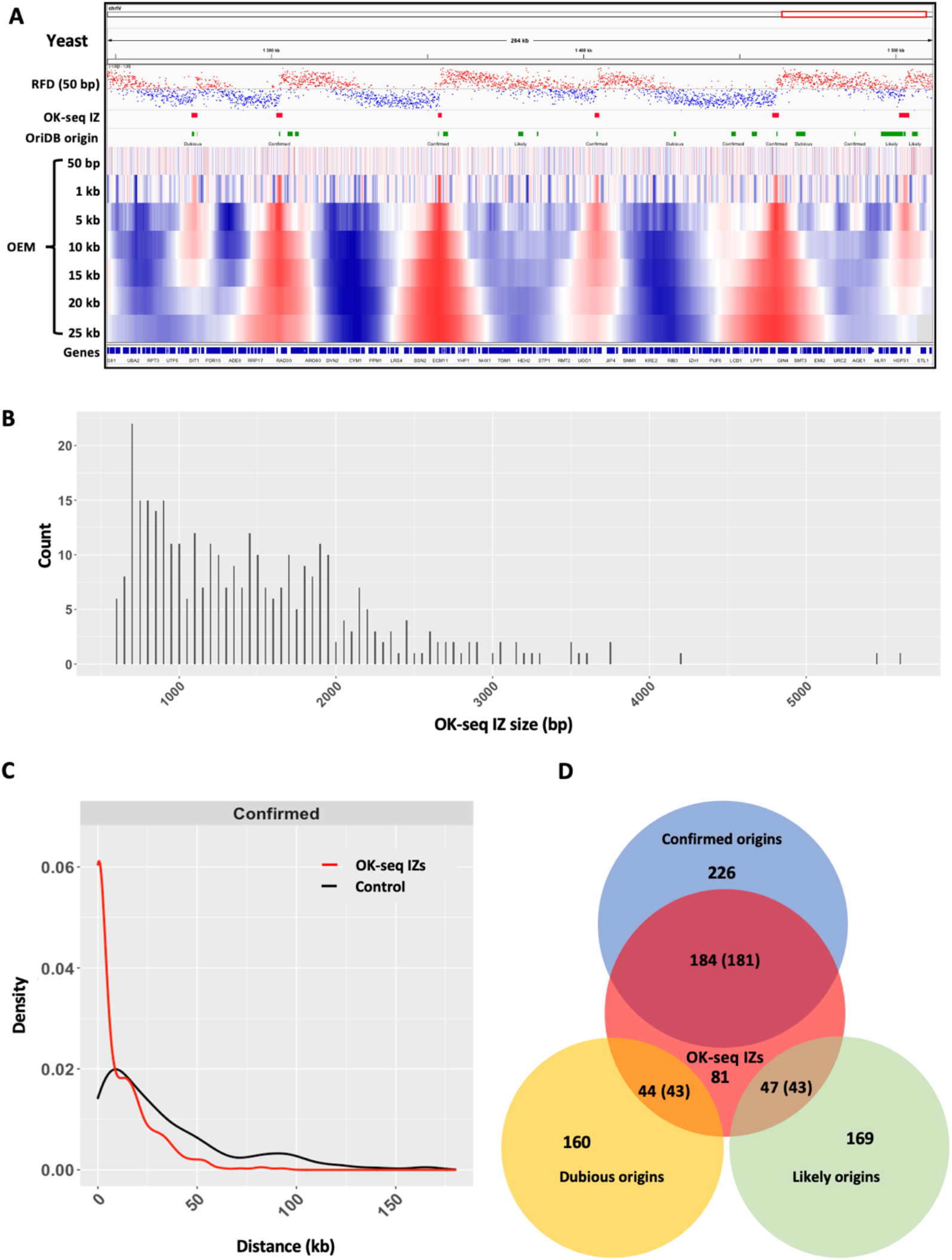
Analysis of OK-seq data by OKseqHMM in Yeast. **(A)** Yeast RFD profile was calculated at 50 bp resolution with the corresponding IZs identified by OKseqHMM, which are highly correlated with the confirmed origins from OriDB (Siow *et al.*, 2012). RFD profile as in Fig. 2C, but with 50 bp resolution. Below, the OEM profiles calculated from 50 bp to 25 kb scales, and the windows with positive and negative OEM values are shown in red and blue, respectively. **(B)** Length distribution of detected OK-seq IZs. **(C)** The density profile shows the distribution (in red) of distances between an IZ detected by OkseqHMM and the closest confirmed origin from OriDB, which is much closer compared with the random simulation control (in black). **(D)** Venn diagram showing the overlap between OK-seq IZs and published origins from OriDB, in which overlap means that the closest distance between each other is less than 1 kb. In case of overlap, the numbers of OK-seq IZ are shown together with the corresponding numbers of OriDB origins indicated in brackets.

### 3.2 Genome-wide replication fork directionality and initiation zone detection for different human cell lines

We then further applied the OKseqHMM to analyze the OK-seq data of human cells. In addition to the published OK-seq data of HeLa MRL2 cells (Petryk *et al.*, 2016), we also generated new additional OK-seq data from HeLa S3 cells, a widely used Encode Tier 2 cell line. The RFD profiles of the two HeLa cell lines are very similar (R=0.86, p<10^−15^), suggesting that a similar replication program and IZ positions are used (Fig. 4A). About 10,000 IZs have been identified in each HeLa cell line (Table 1), two-third of which are common between the two cell lines (Fig. 4B). The conservation of IZs is even higher in the early-replicating regions, with 80% of early IZs being shared between the two HeLa cell lines (Fig. 4B). A very striking difference of human RFD data compared to those of yeast is that, instead of a sharp 1 kb upward transition of RFD at fixed yeast origins, the size of upward transition of RFD, therefore the IZ length, of human cells is around 10-50 kb (average ~30 kb, ~20-folds larger than the IZ of yeast) (Fig. 4A, Table 1). The heatmaps of OEM profiles computed around IZs at different scales show the strongest positive signals at the corresponding scales, i.e., 10 kb scale for the small IZs (<10 kb), 20 kb scale for the IZs of mid-size (20-50 kb) and 50 or 100 kb for the large IZs (>50 kb), respectively (Fig. 4C), confirming that RFD transition is associated with the detected IZ length. This further supports the difference between the yeast and human OK-seq pattern and the accuracy of IZ detection obtained by OKseqHMM.

**Figure 4.**
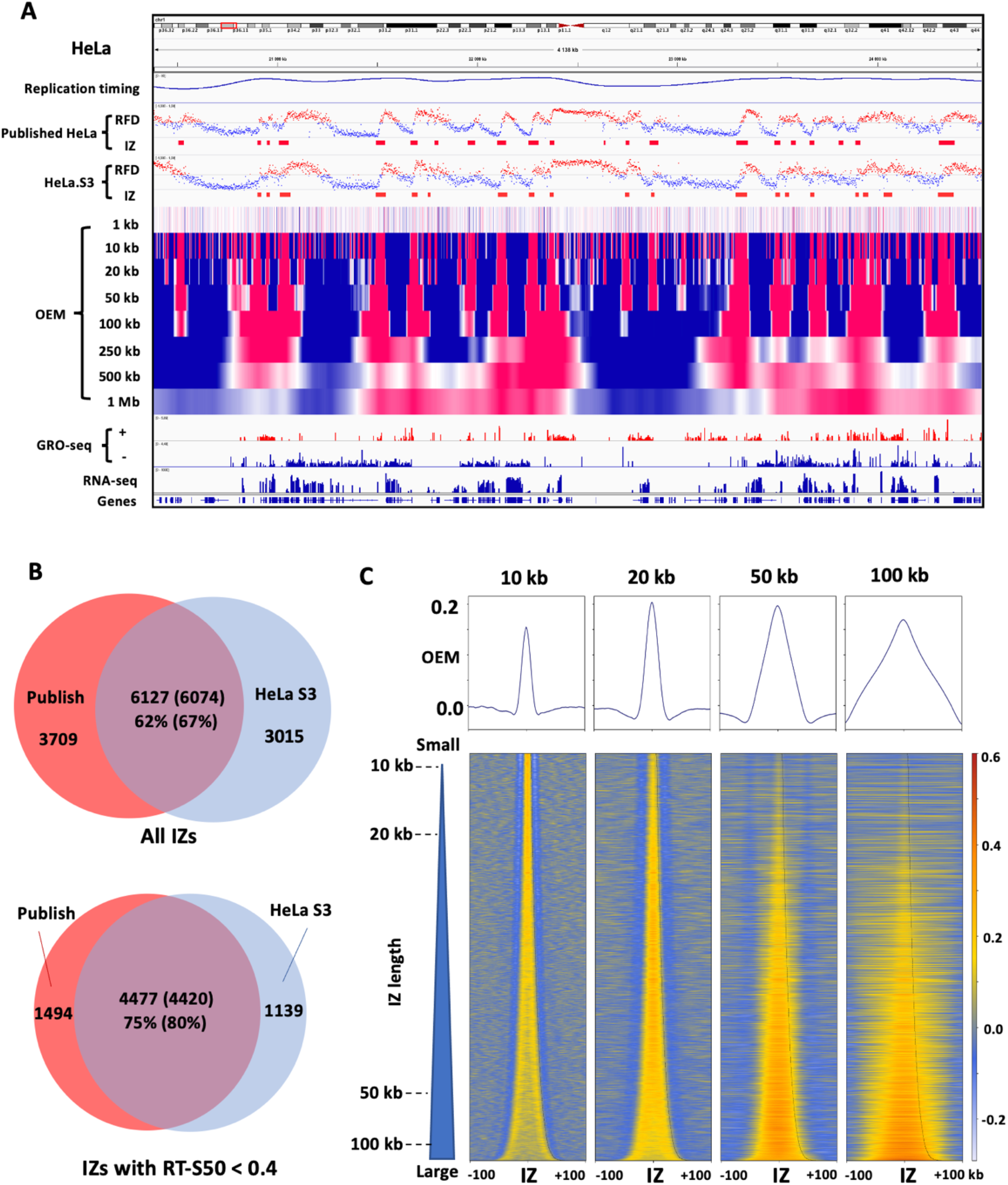
Analysis of OK-seq data of HeLa cells by OKseqHMM. **(A)** Replication timing profile obtained by Repli-seq (Chen *et al.*, 2010), RFD profiles and corresponding detected IZs for published HeLa MRL2 OK-seq data (Petryk *et al.*, 2016) and OK-seq data of HeLa S3 cells generated in the current study, the OEM profiles of HeLa S3 cells from 1 kb to 1 Mb scales, and the transcription data provided by GRO-seq and RNA-seq along a ~4 Mb region on chromosome 1. **(B)** Venn diagrams showing that two-third of Ok-seq IZs matched between the two HeLa cell lines and the overlap goes up to 80 % for the early IZs (with replication timing S50 < 0.4). **(C)** Mean OEM profiles and heatmaps of OEM (heatmap color scale is indicated on the right) around the HeLa S3 IZ centers at indicated scales (i.e., 10, 20, 50 and 100 kb) sorted by the length of detected IZs.

Replication initiation has been previously reported to be enriched within intergenic regions between active genes (Petryk *et al.*, 2016). To demonstrate how our toolkit can help in the analysis of the association between DNA replication and gene transcription, we analyzed the average profiles and the corresponding heatmap of expression level (RNA-seq and GRO-seq) for all detected IZs sorted by their length and confirm that gene transcription presents immediately surround the IZs while with a much lower level within IZs (Fig. 5A). To further compare the distribution of RFD and gene transcription, we calculated the average RFD profile and the corresponding heatmap around TSSs (transcription start sites) and TTSs (transcription termination site) of 16,336 active genes (RPKM > 0) in HeLa cells with an extension ± 50 kb upstream or downstream (Fig. 5B). This clearly indicates a frequent replication initiation (upward transition of RFD) at both regions upstream of TSS and downstream of TTS, which leads to a co-direction between replication and transcription at TSS while a higher head-on TRC at TTS, in agreement with previous publications (Petryk *et al.*, 2016; Promonet *et al.*, 2020).

**Figure 5.**
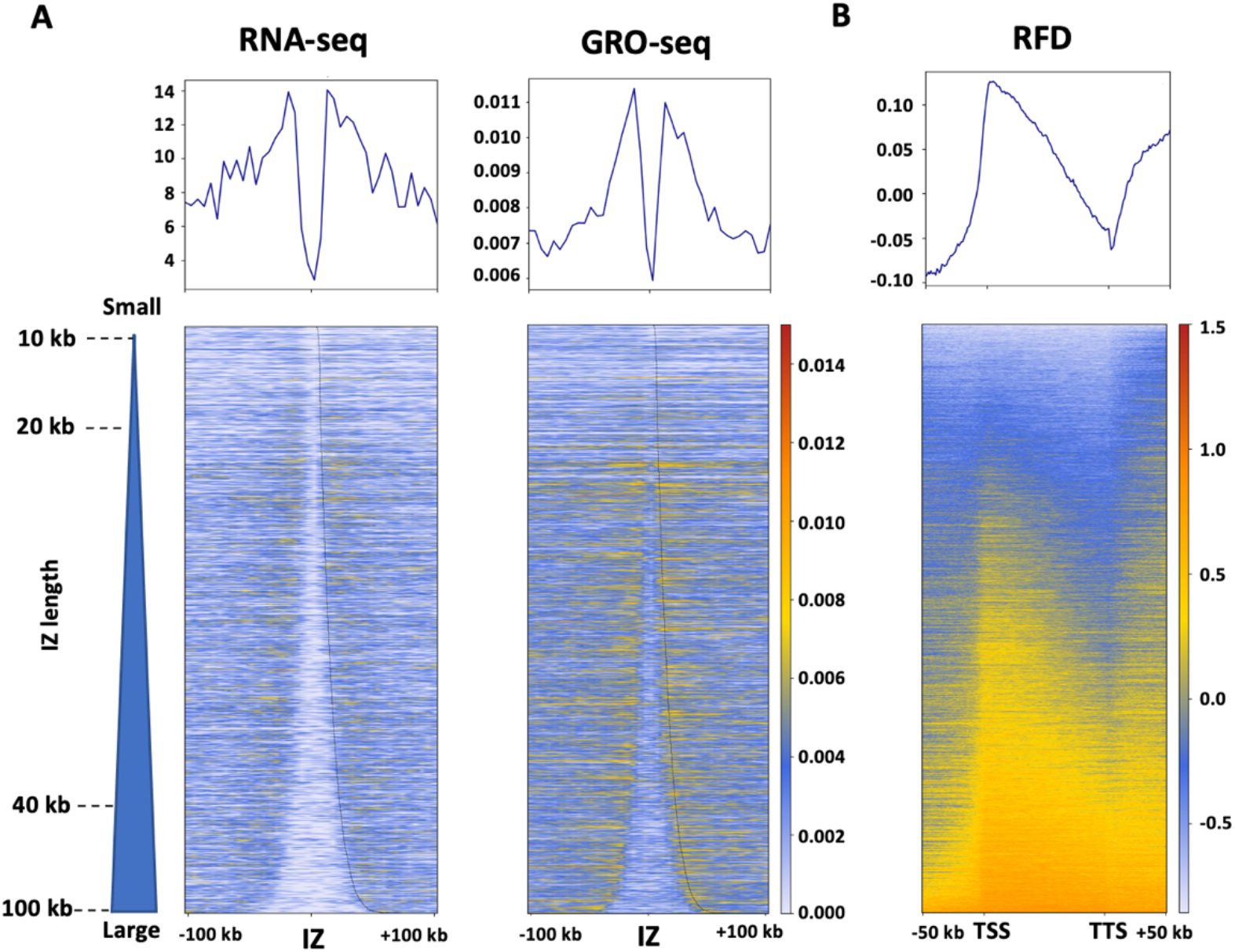
OKseqHMM reveals the coordination between DNA replication and gene transcription. **(A)** Mean profiles and heatmaps of RNA-seq and GRO-seq around the HeLa S3 OK-seq IZ centers. **(B)** Mean profile and heatmap of HeLa S3 RFD between TSS (transcription start site) and TTS (transcription termination site) of active genes with an extension of +/− 50 kb. The heatmap color scales are indicated in each panel.

In addition to OK-seq data of HeLa cell lines, we have gathered and reanalyzed with OKseqHMM, the OK-seq data from previous publications for a large amount of human cell lines of different cell types (Wu *et al.*, 2018; Petryk *et al.*, 2016), such as fibroblast (IMR90), lymphoblastoid (GM06990) and lymphoma (Raji, BL79, IARC385), leiomyosarcoma and leukemia (IB118, TLSE19, K562), and erythroblast (TF1) (Table 1, Fig. 6). OKseqHMM generated high-quality cell-type-specific RFD profiles and robust IZ calling for all data analyzed. The sizes of IZs in different cell types are within the same range (average size between 26 to 36 kb), demonstrating that it is a common feature of human cells.

**Figure 6.**
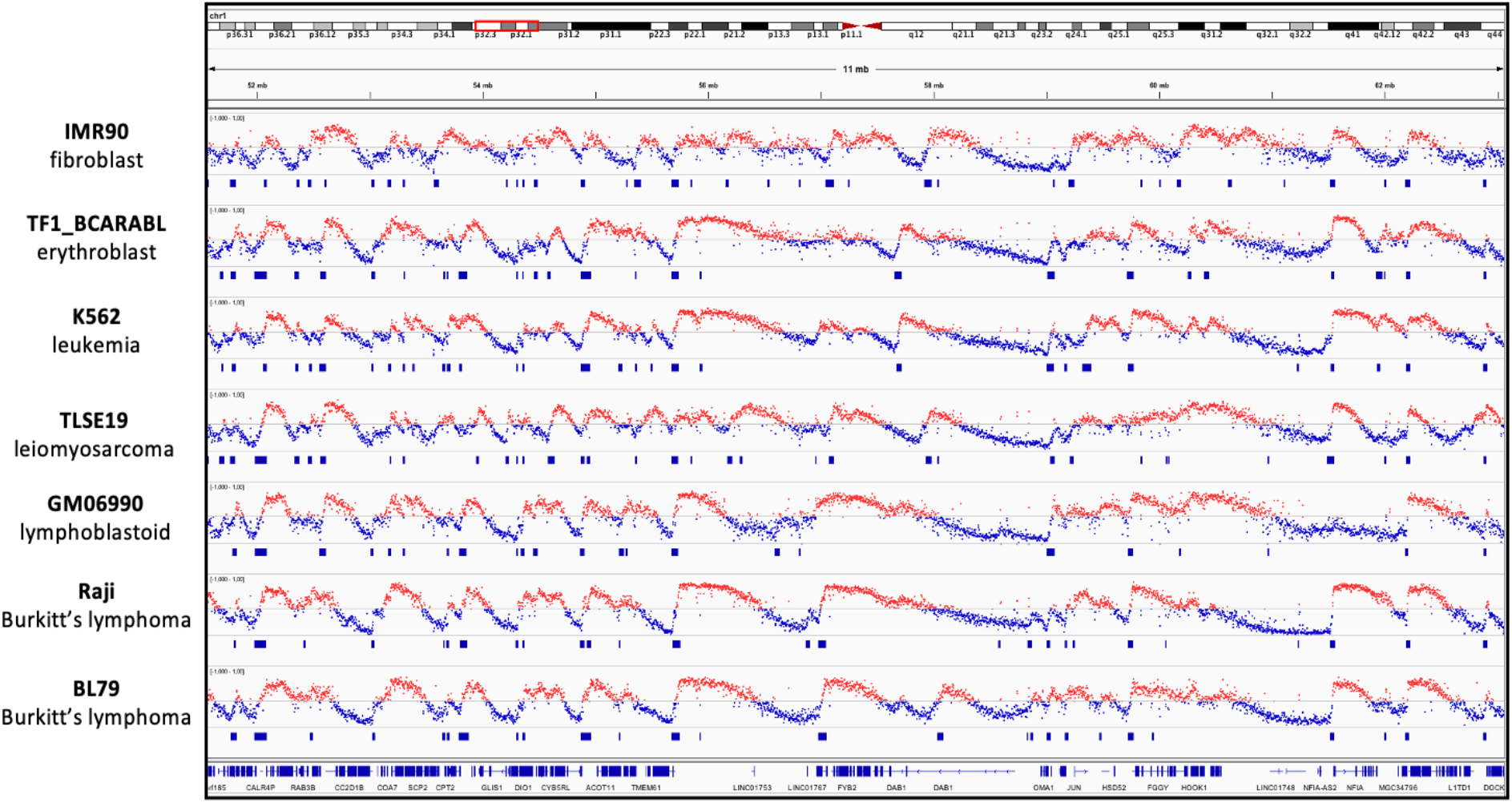
Genome-wide RFD profiles of different human cell lines show the cell-type-specific replication program. Cell-type-specific RFD profiles and the corresponding detected IZs for indicated human cell lines, IMR90, TF1, K562, TLSE19, GM06990, Raji and BL79, respectively.

### 3.3 Extend OKseqHMM to analyze the RFD profiles from other sequencing data

In addition to OK-seq, the OKseqHMM toolkit can be applied to calculate RFD profiles from the sequencing data obtained with other related techniques. As a demonstration, we further extended our toolkit to analyze the published eSPAN (Li *et al.*, 2020) and TrAEL-seq (Kara *et al.*, 2021) data. The RFD data computed from the yeast TrAEL-seq data are very close to those obtained by OK-seq (Fig. 7A, R=0.93, p<10^−15^), and the RFD profile obtained by TrAEL-seq even shows a higher quality with less local noise than the OK-seq RFD profile, which is probably because the TrAEL-seq data used in the analysis contain about two-fold more reads compared with the available OK-seq data. The comparison between the TrAEL-seq IZs, OK-seq IZs and yeast ARSs showed that up to 96% of detected IZs from TrAEL-seq were found within 1 kb distance from a known ARS and around 76% of OK-seq IZs associated with ARSs were also detected by TrAEL-seq (Fig. 7B). We also successfully applied OKseqHMM to the OK-seq and eSPAN data of mouse embryonic stem cells (mESC). However, due to the lower amount of reads for the available dataset of eSPAN data, although we used a larger window size (e.g., 10 kb smoothing window instead of 1 kb window) we still got too noisy RFD profiles to perform a robust IZ calling. Nevertheless, we still obtained a mean RFD profile similar to those of OK-seq around the IZs identified in the mESC OK-seq data (Fig. 7C).

**Figure 7.**
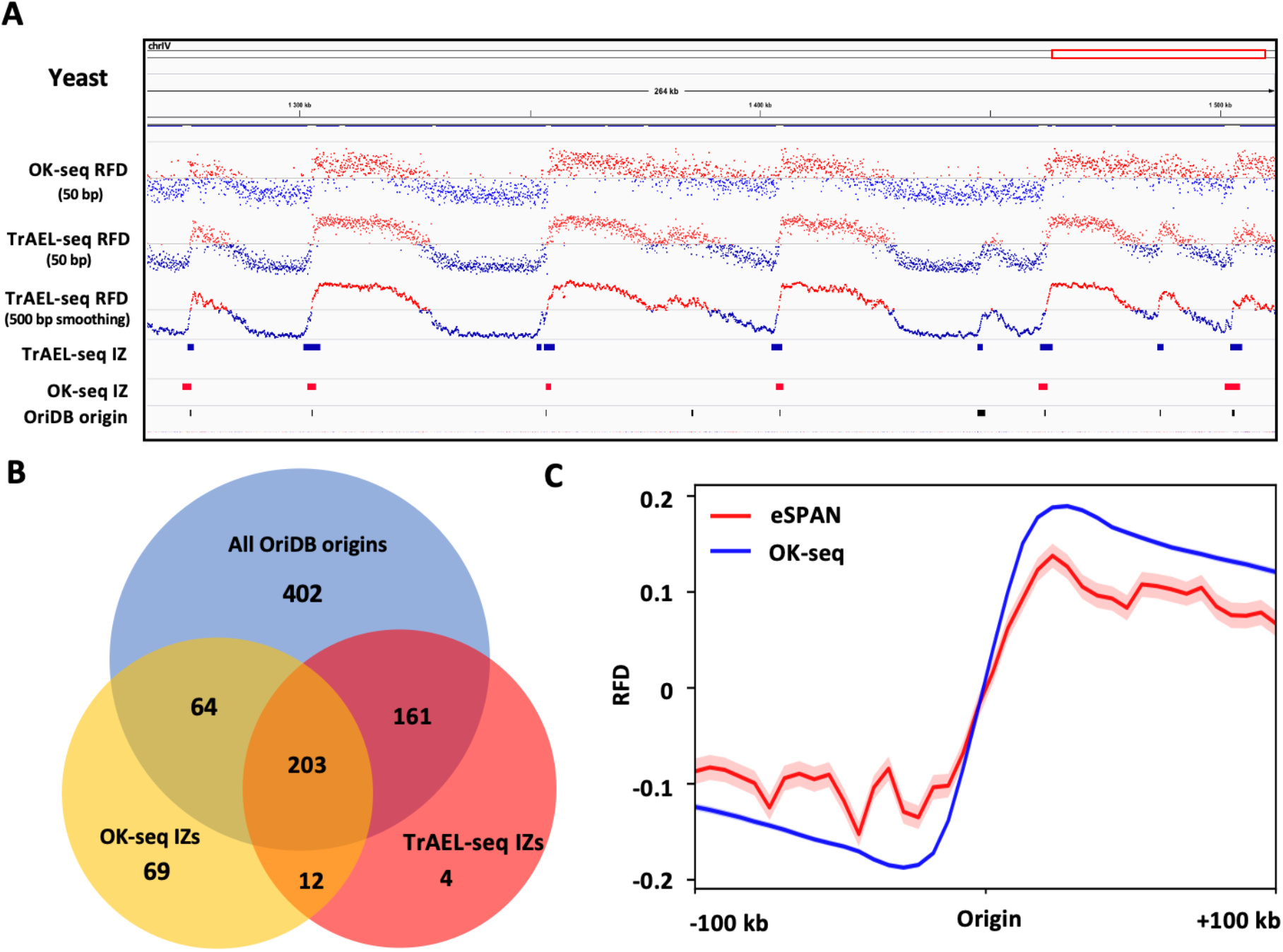
Genome-wide RFD profiles obtained from TrAEL-seq and eSPAN data. **(A)** RFD profiles and the corresponding IZs in 50 bp bin size of OK-seq and TrAEL-seq data of yeast (Kara *et al.*, 2021). The known origins (ARSs) are downloaded from OriDB. **(B)** Venn diagram showing the overlap between OK-seq IZs, TrAEL-seq IZs and published origins (ARSs) from OriDB, in which overlap means that the closest distance between each other is less than 1 kb. **(C)** Metagene average RFD profiles computed from OK-seq of mouse embryonic stem cells (mESC) and H4K20me2 eSPAN data of MCM2-2A mutant cells (Li *et al.*, 2020). Mean and standard error bands are shown for both data, while the standard error bands of OK-seq data are too narrow to be seen.

## 4 Discussion

Genome-wide replication fork directionality data have become an important key in understanding numerous biological processes, such as transcription-replication conflicts, replication-associated mutagenesis, replication couple epigenetic maintenance, etc. Here, we present OKseqHMM, a comprehensive R package, to analyze OK-seq data from various cell types and species to generate and visualize high-resolutive RFD and OEM profiles along the genomes, as well as generate the average profiles/heatmaps on the regions/genes of interest. The toolkit also allows accurate detection of replication initiation/termination zones with an HMM algorithm. To our knowledge, this is the first bioinformatics tool available to date to handle and analyze the RFD data obtained from various techniques.

We successfully applied OKseqHMM to a large amount of available OK-seq data from different species, including yeast, mouse cells as well as numerous normal and cancer human cell lines (Table 1). This provides an important resource for large research communities, who are interested in studying DNA replication programs, transcription-replication conflicts, replication-associated chromatin organization, replication-associated mutations, genome instability and cancer genomics, among others. Importantly, in addition to OK-seq, more and more new techniques have been developed to study DNA replication and are also able to provide the replication fork direction information. These include the methods like eSPAN and SCAR-seq performing stranded sequencing of BrdU or EdU labeled nascent replicated DNA associated with specific histone modifications, or like TrAEL-seq and GLOE-seq based on the single-stranded end presented on specific replicative templates. Here, we demonstrated that OKseqHMM can be applied to analyze data obtained by both kinds of techniques, i.e., eSPAN and TrAEL-seq, and obtained high-quality results (Fig. 7). Notably, techniques like TrAEL-seq, which do not need to incorporate labels and need fewer cells to generate a high-quality RFD profile compared to OK-seq, will provide a good alternative to study DNA replication and genomic instability in different cell types within various stress conditions.

It should be noted that the initiation parameters, such as the transition and emission probabilities, are defined based on the OK-seq datasets of human cells. Although we have shown in the current study that they are quite robust and can be also applied to OK-seq data of yeast (Fig. 1) and mouse cells (Fig. 7, Table 1) to obtain satisfactory results, they might need to be adjusted based on the sequencing-depth and data quality of other datasets, in order to have an optimal IZ/TZ calling. In the future, with technical improvement, we might be able to further extend the OKseqHMM to study the extrinsic (cell-to-cell) or intrinsic (homolog-to-homolog) variability of DNA replication, if we can further extend the relative techniques to obtain data at the single-cell level and/or in an allele-specific manner as recently achieved for the replication timing study (Dileep and Gilbert, 2018; Gnan *et al.*, 2021).

## Data Availability

The bioinformatics tool and all processing data underlying this article are available at the GitHub page of the team https://github.com/CL-CHEN-Lab/OK-Seq. And the raw sequencing data of OK-seq are available with the corresponding accession numbers indicated in Table 1. The OK-seq data of HeLa S3 cells were generated as described in (Petryk *et al.*, 2016) and are available at GEO. The RNA-seq and GRO-seq data of HeLa cells are from (Promonet *et al.*, 2020) and (Andersson *et al.*, 2014), respectively. The known yeast origins (ARSs) are downloaded from OriDB (http://cerevisiae.oridb.org/) (Siow *et al.*, 2012).

## Acknowledgements and funding information.

This work was supported by the YPI program of I. Curie; the ATIP-Avenir program from Centre national de la recherche scientifique (CNRS) and Plan Cancer [grant number ATIP/AVENIR: N°18CT014-00]; the Agence Nationale pour la Recherche (ANR) [grant number ReDeFINe - 19-CE12-0016-02, TELOCHROM - 19-CE12-0020-02]; and Institut National du Cancer (INCa) [grant number PLBIO19-076]. YL thanks ANR for providing her Ph.D. fellowship. The authors would like to thank Joseph Josephides for critical reading, and to acknowledge the high-throughput sequencing facility of Institute for Integrative Biology of the Cell (I2BC) for its sequencing expertise. We declare no competing interests and no potential conflicts of interest.

